# Single cell guided deconvolution of bulk transcriptomics recapitulates differentiation stages of acute myeloid leukemia and predicts drug response

**DOI:** 10.1101/2022.12.09.519738

**Authors:** E Onur Karakaslar, Jeppe Severens, Elena Sánchez-López, Peter A van Veelen, Mihaela Zlei, Jacques JM van Dongen, Annemarie M. Otte, Constantijn JM Halkes, Peter van Balen, Hendrik Veelken, Marcel JT Reinders, Marieke Griffioen, Erik B van den Akker

## Abstract

The diagnostic spectrum for AML patients is increasingly based on genetic abnormalities due to their prognostic and predictive value. However, information on the AML blast phenotype regarding their maturational arrest has started to regain importance due to its predictive power on drug responses. Here, we deconvolute 1350 bulk RNA-seq samples from five independent AML cohorts on a single-cell healthy BM reference and demonstrate that the morphological differentiation stage (FAB classification) could be faithfully reconstituted using estimated cell compositions (ECCs). Moreover, we show that the ECCs reliably predict *ex-vivo* drug resistances as demonstrated for Venetoclax, a *BCL-2* inhibitor, resistance specifically in AML with CD14+ monocyte phenotype. We further validate these predictions using in-house proteomics data by showing that *BCL-2* protein abundance is split into two distinct clusters for NPM1-mutated AML at the extremes of CD14+ monocyte percentages, which could be crucial for the Venetoclax dosing for these patients. Our results suggest that Venetoclax resistance predictions can also be extended to AML without recurrent genetic abnormalities (NOS), and possibly to MDS-related AML and secondary AML. Collectively, we propose a framework for allowing a joint mutation and maturation stage modeling that could be used as a blueprint for testing sensitivity for new agents across the various subtypes of AML.

## Introduction

Acute myeloid leukemia (AML) is an aggressive hematological cancer of the myeloid lineage. AML is caused by a combination of relatively few genetic alterations that are predominantly somatically acquired and cooperatively induce a maturation arrest in combination with rapid uncontrolled proliferation of immature myeloid precursor cells. The prognosis of AML is highly dependent on the presence of such recurrent genetic alterations and varies from > 90% cure rates to less than 10%^1^. Therefore, the current WHO classification primarily defines AML subtypes according to the presence of eleven recurrent genetic aberrations (RGA) changes and an added heterogeneous umbrella subtype composed of a highly diverse set of relatively rare RGAs ^3–6^. Only AML cases that lack any detected RGA are characterized according to their maturation stage according to the French-American-British (FAB) classification ^2^.

Recently, the maturation and differentiation stage of AML blasts has gained importance due to a striking association with sensitivity or resistance to new drugs ^7–9^. AML differentiation stage can be assessed by diagnostic flow cytometry more objectively than by morphological examination alone. However, flow cytometry is limited to a relatively low number of differentiation markers. Gene expression profiling by bulk RNA-sequencing (RNA-seq) is an attractive alternative technique as it allows both calling of genetic aberrations and estimation of cell subsets, i.e. estimated cell composition (ECC) from the same sample^10–12^. Reported attempts to estimate ECCs by deconvoluting bulk AML samples utilizing single-cell RNA-seq as an *in-silico* reference, were mostly focused on detection of survival differences without performing thorough validation of the ECCs and/or used leukemic samples as reference^13–15^, which prevents assessing whether ECCs are tissue- or sample-specific.

Here, we perform deconvolution of 1350 AML transcriptomic samples via a healthy single-cell reference while validating our findings with our in-house flow cytometry data. Of note, we demonstrate that the ECCs recapitulate the entire FAB landscape (M0-M7). Then, using these ECCs we predict *ex-vivo* drug resistance data from literature and show the agreement of these results at protein level with the help of in-house proteomics data in AML patients, for whom we also had acquired gene expression data. To conclude, we hereby propose a transcription-based single-cell guided deconvolution framework to assess the drug effectiveness to different maturational stages of AML. We also provide our framework as a CRAN R package available at https://github.com/eonurk/seAMLess.

## Results

### Deconvolution pipeline recapitulates healthy and malignant hematopoiesis

We initially created a healthy bone marrow (BM) reference atlas from single-cell transcriptomics data (**Figure 1A**). For this purpose, we integrated data from 69,101 cells covering 439 genes from three publicly available datasets (two full-transcriptome and one targeted) from two studies^16,17^(**Figure 1B**,Methods). As expected, T-cells and B-cells formed separate clusters in the UMAP plot, whereas the myeloid lineage cells clustered together. To differentiate early myelopoiesis, we distinguished more than 3,349 hematopoietic stem cells (HSC) and 1,432 erythro-myeloid progenitors (EMP), 2,939 lymphoid multipotent progenitors (LMPP), as well as 3,508 granulocytes-monocytes progenitors (GMP) (**Figure 1C**). Inter-individual variability, possibly due to age differences, mostly affected T- and B-cells and did not influence overall clustering (**Figure S1A**). Homogeneous distribution of the studies on the UMAP plot showed successful integration of the different datasets (**Figure S1B**).

**Figure 1.**
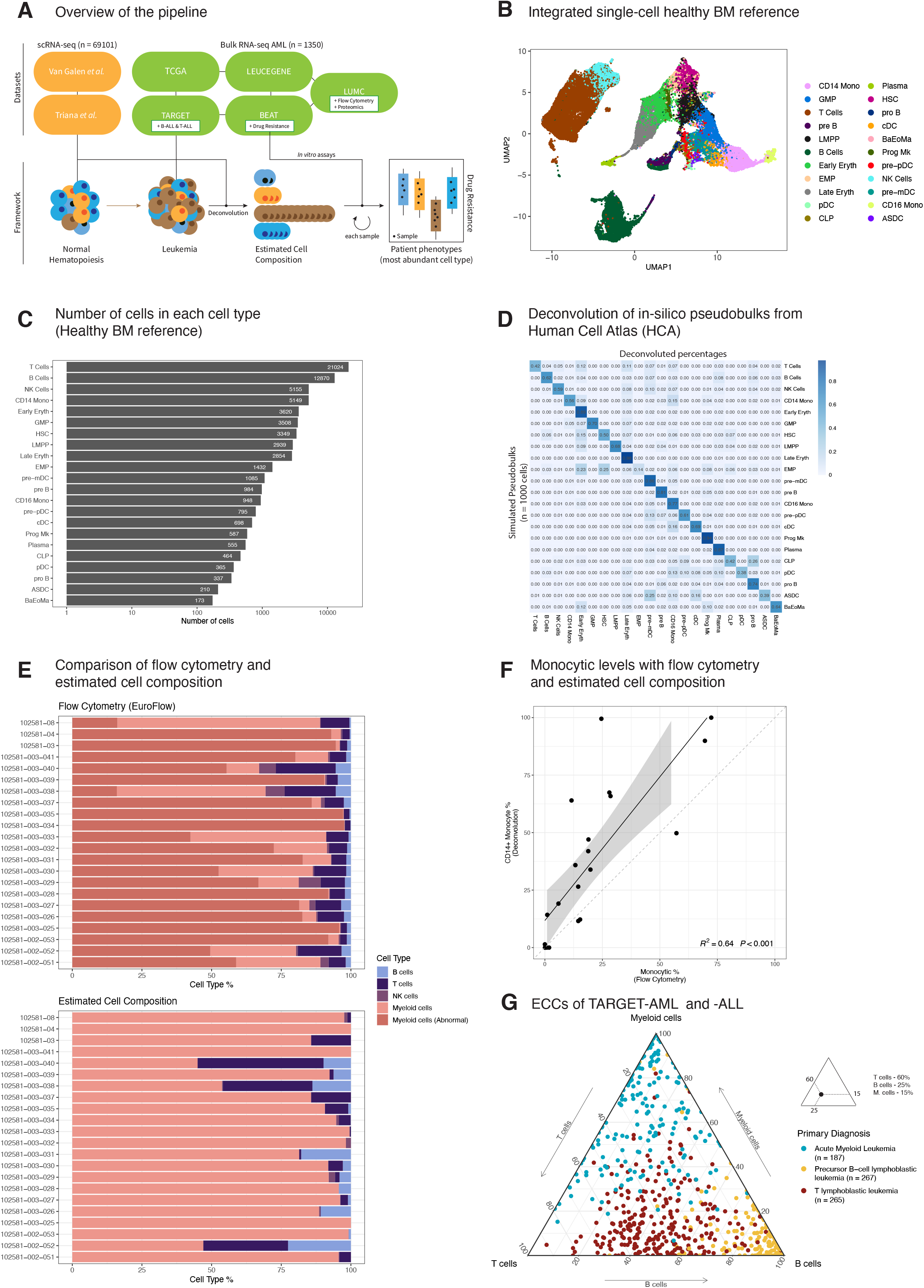
Schematic of the study and compositional validation. **A** The overview of the study. Integrated cells for healthy bone marrow (BM) dataset were collected from two studies^16,17^. Five independent bulk transcriptomic AML cohort (TCGA-LAML, BEAT-AML, TARGET-AML, LEUCEGENE and LUMC) samples (n=1,350) were deconvoluted. Drug resistance data (n=122) from BEAT-AML were predicted using ECCs. Additional ALL samples (n=532) from TARGET study were used for validating deconvolution framework. Proteomics (n=39) and flow cytometry samples (n=22) from LUMC cohort were used for further validation of the framework. **B** UMAP plot of integrated healthy BM (n=69,101) cells. Annotations are lifted via Azimuth framework. T and B cell subset were merged, resulting into 22 cell types (abbreviations; GMP: Granulocyte Monocyte Progenitors, LMPP: Lymphoid Primed Multipotent Progenitors, NK: Natural Killer, EMP: Erythroid Megakaryocyte Progenitor, pDC: Plasmacytoid Dendritic Cell, CLP: Common Lymphoid Progenitor, HSC: Hematopoietic Stem Cell, BaEoMA: Basophil Eosinophil Mast Progenitor, Prog Mk: Progenitor Megakaryocyte, pre-mDC: Precursor Myeloid Dendritic Cell, ASDC: AXL+ Dendritic Cell). **C** Barplot for cells per cell type of the healthy BM. X-axis is in log10 scale. **D** Heatmap showing the fraction of deconvoluted cell types for simulated pseudobulks from HCA subset. Each row indicates a simulated pseudobulk with an overabundant cell type (80%) and adds up to 1, and each column is a cell type from the healthy BM reference. **E** Cellular composition of the same samples via flow gating of orientation tube (ALOT) and ECCs (RNA-seq). Abnormal assignment within myeloid cells is determined by flow gating, thus only exists for flow cytometry. **F** Percentage of monocytic subsets measured by flow cytometry and ECCs. **G** Ternary plot showing the ECCs of TARGET AML and ALL cohorts (n=719), each dot represents a sample and corners indicate a major cell type and colors indicate the primary diagnosis of each leukemic sample (Supplementary Table S1).

We next performed *in silico* experiments to validate our deconvolution set-up. We first used the healthy BM reference to create *in silico* mixed bulk samples with known cell compositions. Pseudobulk profiles were simulated with one abundant cell type (80% of the cells) with the remaining cells being a mix created by random selection (Methods). Then, MuSiC^12^ with the default settings was used to deconvolute the simulated pseudobulk profiles into their respective cell types (**Figure S1C**). To validate on an independent dataset, we also simulated pseudobulks from 40,000 healthy bone marrow cells of the human cell atlas (HCA)^18^(**Figure S1D**) and performed deconvolution via MuSiC. All simulated pseudobulk profiles were successfully deconvoluted into their respective cell types (matching to the most abundant cell type) apart from EMP (**Figure 1D**). For this profile, the annotations were shared mostly among EMP (14%) as well as HSC (25%) and Early Erythrocytes (23%). This discrepancy might be explained by the transcriptional similarity of these cell types (**Figure 1B**).

Next, we analyzed our in-house diagnostic flow cytometry data (EuroFlow^19^ panels; see Methods) for 22 AML samples with matched bulk RNA-seq data. An overview of mean fluorescence intensity (MFI) values for these samples’ abnormal cells after staining with antibodies for 33 markers distributed over 7 tubes (1 Orientation + 6 AML assignment tubes) is shown in **Figure S1E** (Supplementary Table S1). In line with our expectations, the most abundant cell types for all samples were in the myeloid lineage for both flow cytometry analyses and estimated cell compositions (ECCs) (**Figure 1E**, Supplementary Table S4). To investigate whether monocytic AML can be accurately distinguished from AML with more stem cell-like phenotypes, we plotted CD14+ monocyte percentages as determined by deconvolution against MFI values of antibodies of monocytic markers (CD11b, CD64, IREM2, and CD14) on all BM cells without gating, and observed statistically significant correlations for 3 markers (CD11b, CD64, IREM2), with CD64b being most significant (R^2^= 0.43, P < 0.001) (**Figure S1F**). Also, percentages of AML cells assigned to the monocytic subset by EuroFlow panels and ECCs showed statistically significant correlations (R^2^= 0.64, P< 0.001) (**Figure 1F**).

Lastly, we downloaded publicly available TARGET AML and ALL (B-ALL and T-ALL) data and deconvoluted these samples (n=719) with the healthy BM reference (**Figure 1G**, Supplementary Table S1) to show that different leukemic phenotypes could be captured by deconvolution. The estimates of cell type abundances in the three immune lineages (myeloid, B- and T-lymphoid) were visualized using a ternary plot. As expected, each type of acute leukemia was closer to their respective cell type of origin (ECCs), most compact being B-ALL (yellow samples on the right corner), showing the ECCs’ ability to capture the patients’ immune phenotypes at major cell type levels. Together, these benchmarking and validation analyses demonstrated that given a healthy single cell BM transcriptomic atlas, the cell type proportions can be recapitulated faithfully via deconvolution from bulk transcriptomic leukemia cases.

### Deconvolution of bulk AML transcriptomics reveals the dominating immune fraction

To investigate heterogeneity in cell composition of AML, we next applied our framework to deconvolute five independent bulk transcriptomic AML studies, i.e. TCGA-LAML^20^ (n=151), BEAT-AML^21^ (n=460), TARGET-AML^22^ (n=187), LEUCEGENE^23^ (n=452) and our cohort LUMC^10^ (n=100) totaling in 1350 samples from 1,267 patients (**Figure S2A**). The results are shown in **Figure 2A** as a heatmap where each sample was decomposed into the 22 cell types from the healthy BM reference. We also added information on patients’ clinical blast counts, FAB^2^, WHO 2016^4^, and ELN 2017^6^ classes for an overarching picture of AML landscape, and we also calculated the stemness score^24^, which is a gene expression signature for patient prognosis trained on engraftment capacity of AML in immunodeficient mice (Supplementary Table S2). Each sample was assigned to the most abundant cell type as determined by deconvolution (AML phenotype) and arranged according to their fractions within each phenotype. The data showed a clear dominance of one immune cell type for most of the cases, clearly indicating that maturational arrests and lineage skewing are leukemic properties that can be readily assessed using transcriptome sequencing. The majority of AML cases was typically estimated to be dominated by myeloid cells (**Figure 2A, S2A, S2B, S3D**). As notable exceptions, a few pediatric AML samples from the TARGET cohort showed ECC profiles dominated by T-cells or B-cells. This effect is most likely caused by prominent infiltration of the bone marrow by these types of lymphocytes besides the AML blast population and therefore precludes the assignment of the AML blasts to a distinct myeloid cell type. As previously reported these cases can nevertheless be considered as a special subgroup of pediatric AML, and specifically T-cell dominated patients were recognized for poor survival^25^. Furthermore, in line with previous reports, patients with acute promyelocytic leukemia (APL), classified as AML-M3 by FAB, were correctly assigned to have a cell composition dominated by granulocyte-monocyte progenitors (GMPs). Besides APL, there are also AML cases dominated by GMP and large groups of AML assigned as monocytic AML or AML with an earlier HSC or EMP phenotype, again demonstrating that bulk transcriptomics can be used to capture the stage of arrest of AML during hematopoiesis by deconvolution.

**Figure 2.**
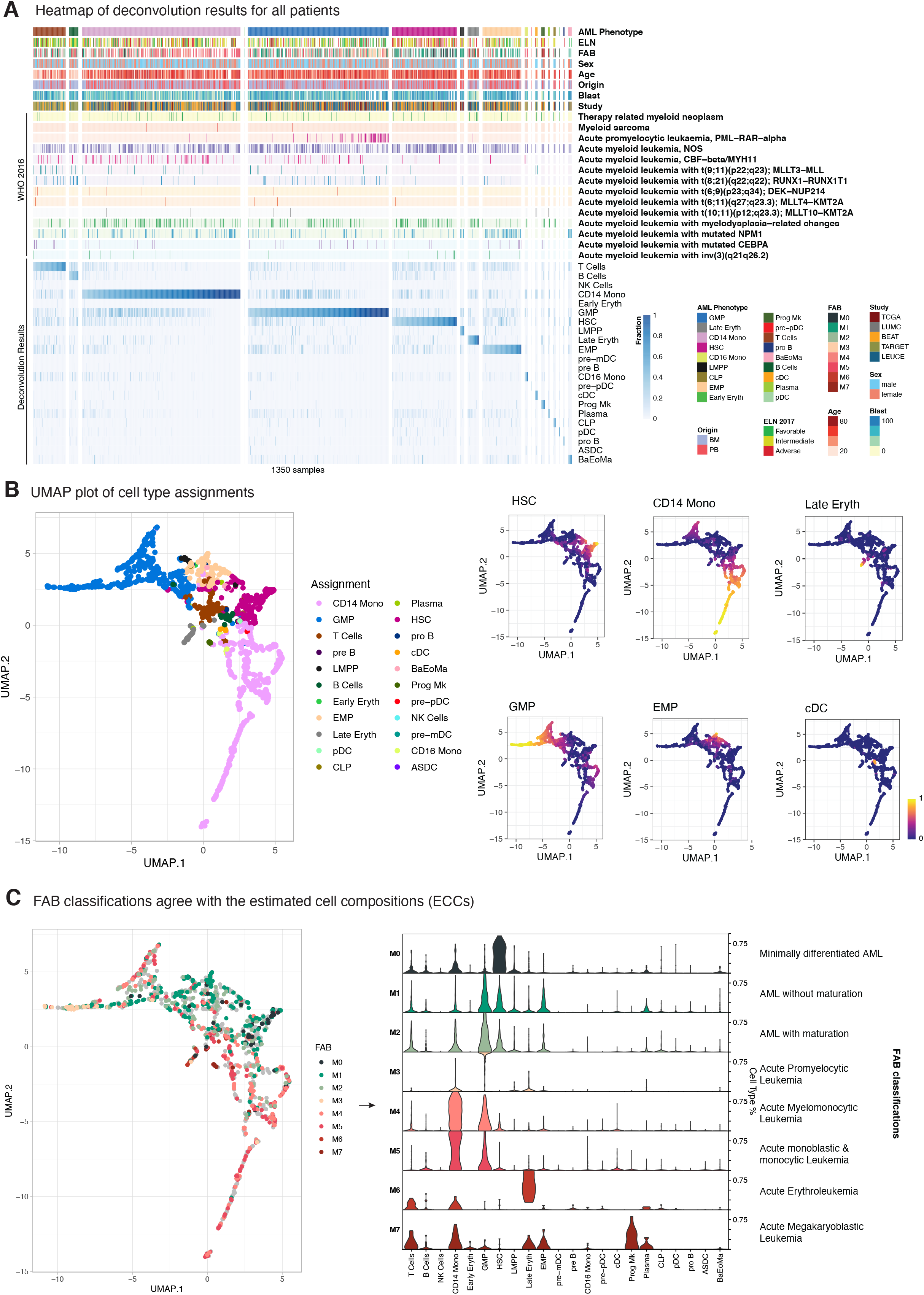
Capturing differentiation stages with ECCs. **A** Heatmap showing the deconvoluted fractions for of five AML cohorts (TCGA-LAML, BEAT-AML, TARGET-AML, LEUCEGENE and LUMC). Each column represents a sample, which is deconvoluted into 22 cell types (bottom part with blue) via healthy single cell BM reference. Top part of the annotations shows the provided meta data (FAB, WHO and ELN classifications, origin, sex and reported blast percentages) for these cohorts. Each patient was assigned to its most abundant deconvoluted cell type (AML phenotype) and samples within each assignment were sorted according to the assigned phenotype. _B_ UMAP plot of deconvolution levels annotated via most abundant cell type as the left panel; and on the right, ECCs of 6 out of 22 cell types were shown in continuous scale (HSC, CD14+ Monocytes, Late Erythrocytes, GMP, EMP and cDC). **C** Similar to UMAP plot in (B) but colored with FAB classifications (left panel), and the fractions of cell types for each of these FAB classifications (right panel).

### Deconvolution of bulk AML transcriptomics agrees upon stemness score

To comprehensively visualize changes in ECCs in relation to metadata, ECCs of AML samples were visualized using a UMAP plot. As expected, samples with similar ECCs clustered, as shown after annotating the samples for their most abundant cell type (**Figure 2B** – left panel). When superimposing the deconvoluted percentages of 6 major cell types on the same UMAP plot (**Figure 2B** – right panels), gradual shifts in cell composition became apparent, in particular, APL samples populated the extreme extension of the GMP cluster. Moreover, a gradient towards monocytic outgrowth and conversely to a high stemness was also observed. Comparisons with the clinical blast percentage and previously published stemness score showed an overall agreement with ECCs, HSCs and EMPs having a higher score than CD14+ Monocytes and GMPs (**Figure S3A, S3B**). Late Erythrocyte also had high stemness score, albeit with large variation. By dividing AML into cases with high and low stemness scores, we observed similar compositional changes with abundance of HSCs and EMPs in AML with high stemness scores in all cohorts. Late erythrocytes, however, were not consistently enriched in all cohorts. Furthermore, we noticed that in TARGET, T cells are abundant in AML with high stemness scores (**Figure S3C**). This raises the question whether the stemness score, which has been trained on adult AML, is useful to predict prognosis of pediatric patients particularly considering that large subtypes of adult AML such as APL and AML with mutated NPM1 are rare in pediatric AML. The consistent enrichment of HSCs and EMPs in AML with high stemness scores in all cohorts, however, further supports correct determination of the cell composition of AML by deconvoluting bulk transcriptomics.

### Estimated cell composition recapitulates FAB classes

Clear associations were also observed between cell type assignments by ECC and FAB classification status (left panel of **Figure 2C**). Furthermore, distinct distributions of cell type-defining gene expression profiles became evident for each FAB AML type by plotting continuous values of deconvolutions rather than categorical assignments: Mo (minimally differentiated AML) cases had high levels of the HSC-defining signature, whereas these levels decreased and EMP- and GMP signatures appeared in M1 (AML without maturation) and M2 (AML with maturation). As expected, M3 (APL) cases were almost completely dominated by the GMP signature, while M4 (Acute Myelomonocytic Leukemia) and M5 (Acute monoblastic/monocytic leukemia) samples resembled CD14+ monocyte cells and GMP. The small groups of M6 (Acute Erythroleukemia) and M7 (Acute Megakaryoblastic Leukemia) AML were dominated by the signatures of Late Erythrocytes and Megakaryocyte Progenitors, respectively. These data demonstrated and confirmed that our single cell guided deconvolution strategy successfully captures the maturational arrest of AML cells at different differentiation stages of hematopoiesis.

### Estimated cell composition captures genetic subtype-specific resistances to various drugs

To explore whether ECCs convey information on drug resistances, ex *vivo* drug response data of 122 small molecule inhibitors provided as area under the curves (AUC) for 363 AML samples from BEAT were downloaded. A higher AUC indicates that cancer cells are relatively resistant since higher drug concentrations are needed to induce cell death. To predict drug resistance for AML samples with calculated ECCs, we trained random forest (RF) models per drug via leave-one-out cross validation (LOOCV) setting and then calculated the Spearman ρ values of these models’ predictions (**Figure S4A;** Supplementary Table S5-S7). We also stratified these results according to WHO classifications (**Figure 3A, S4B**, Supplementary Table S8).

**Figure 3.**
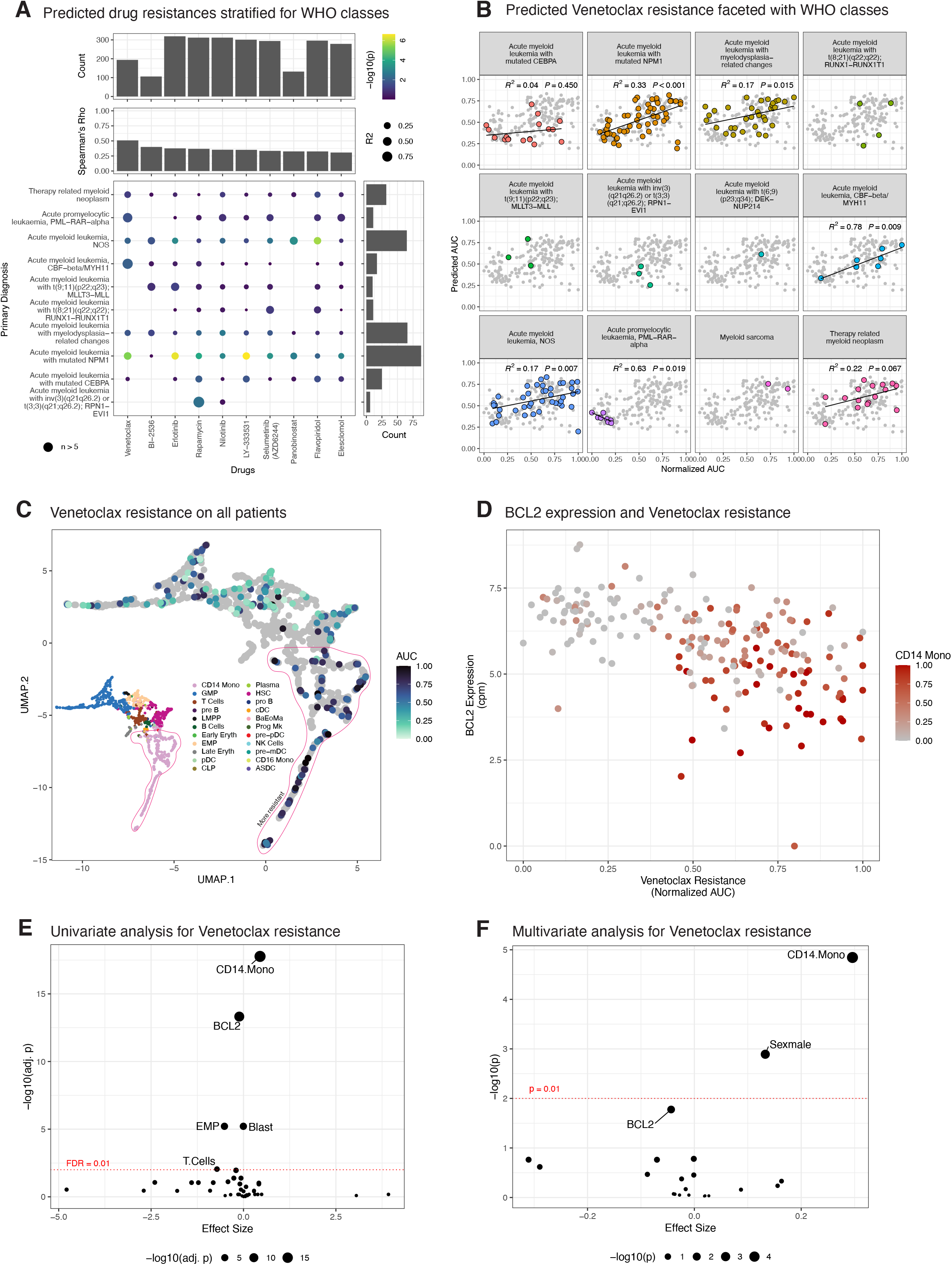
*Ex-vivo* drug response predictions. **A** Correlation of predicted AUC and normalized AUC of *ex-vivo* drug resistances split by WHO classes (top 10 drugs). Bigger bubble corresponds to higher correlation, and colors indicate statistical significance. Only groups with >5 samples were fit linear models thus R^2^ and p values are shown. **B** Random Forest predictions of Venetoclax resistance via ECCs with LOOCV (ρ = 0.509; Supplementary Table - BEAT AUC RF Predictions). Each dot represents a patient. Higher resistance (AUC) means more cells are viable after administering the drug. **C** Samples with normalized Venetoclax resistance data are shown on the UMAP plot of ECCs, along with phenotype map from (**Figure 2B**). Patients without available data were shown as gray dots for the sake of completeness. **D** Venetoclax resistance vs *BCL-2* expression; also, CD14+ Monocyte percentages for these patients were annotated. Expression data was count-per-million (cpm) normalized. **E** Univariate analyses of different attributes (i.e., CD14+ Monocyte %, reported blast percentages etc.) vs normalized Venetoclax resistance. P values were corrected using Benjamini-Hochberg procedure and the names of the attributes are shown for FDR > 0.01. **F** Similar to **E; A** multivariate model is associated with different attributes, and the names of the attributes with P> 0.01 is shown. Full list of attributes’ p values and estimates are provided in Supplementary Table for both **E** and **F**.

Drug responses to the EGF-R inhibitor Erlotinib (Spearman ρ = 0.376 and the mTOR pathway inhibitor Rapamycin (Spearman ρ = 0.368) showed high correlations with AML with mutated NPM1 (n = 77, R^2^ = 0.29, P= 3.7×10^7^) and inv(3) (n = 6, R^2^= 0.91, P= 3.2×10^3^), respectively. Notably, responses to Flavopiridol (Spearman ρ = 0.325), a CDK kinase inhibitor, showed strong correlations with ECC within the group of AML-NOS (n = 77, R^2^= 0.31, P= 3.7×10^7^) while showing higher resistance towards AML with more stem cell like cell phenotypes (**Figure 3A, S5A, S5B**). The strongest correlation (Spearman ρ = 0.509), however, was observed for Venetoclax (ABT-199), a BCL2 inhibitor (**Figure S5C**). Annotation with primary diagnosis from WHO classes (**Figure 3B**) revealed that PML-RARA-carrying APL samples (n = 8) are dominated by GMPs are sensitive for Venetoclax, whereas AML with CBFB-MYH11 (n = 7) had the best fit (R^2^= 0.78, P = 0.009) for Venetoclax resistance, although the small sample size of CBFB-MYH11 cases prohibits drawing robust conclusions. Larger groups of AML, however, such as NPM1 mutated cases (n = 55, R^2^= 0.33, P < 0.001), and AML-NOS (n = 42, R^2^= 0.17, P = 0.007), together accounting for approximately 58% of AML, also showed a clear trend for Venetoclax resistance (**Figure 3B**). Annotating drug responses with predicted resistances by deconvolution (**Figure S5C**)revealed that CD14+ Monocytic AML are relatively resistant to Venetoclax. This resistance of monocytic AML was also clear by overlaying the AUC for Venetoclax onto the UMAP plot (**Figure 3C**) or boxplots (**Figure S5D**) (one-way ANOVA p = 6.31×10^13^; Supplementary Table S9). Also, within the group of NPM1 mutated samples, which is the largest class of AML, CD14+ Monocyte dominated cases were most resistant to Venetoclax (**Figure S5E, S5F;** one-way ANOVA p = 7.6×10^4^; Supplementary Table S10). Briefly, these findings suggest that information on the ECCs of AML might yield therapeutic implications even within one genetic subtype.

### Estimated CD14+ Monocyte percentages predict Venetoclax resistance better than *BCL-2* mRNA expression

Since Venetoclax is targeting the anti-apoptotic *BCL-2* protein, we next checked *BCL-2* mRNA expression levels in AML and overlaid the CD14+ Monocyte percentages for these cases (**Figure 3D**). The data showed that low BCL2 expression indeed correlated with strong resistance to Venetoclax. However, a few samples were resistant despite relatively high *BCL-2* expression (samples at upper right corner in **Figure 3D**). As the majority of these cases had a CD14+ Monocyte phenotype, we univariately associated *BCL-2* expression and CD14+ Monocyte percentage to investigate which factor best explains the Venetoclax response (**Figure 3E**).

We also associated metadata such as reported blast percentages, ELN status and primary diagnosis to determine their effect sizes and significance (Supplementary Table S11). This analysis demonstrated that both CD14+ Monocyte percentages (adjusted P = 1.7×10^18^) and *BCL-2* mRNA expression (adjusted P = 4.8×10^14^) were associated significantly with the Venetoclax response. Next, we created a multivariate model to investigate whether the significance of CD14+ Monocyte percentages diminishes along with the presence of *BCL-2* expression in the same model (**Figure 3F**, Multivariate Venetoclax Tab in Supplementary Table 12). In this model, CD14+ Monocyte percentages remained significantly associated with Venetoclax resistance (P = 1.92×10^5^), while *BCL-2* expression was below the significance threshold of 0.01 (P = 0.015). In conclusion, these results indicate that cellular composition is a more robust marker than *BCL-2* mRNA expression to predict Venetoclax resistance, specifically for AML from NOS and NPM1 mutated patients.

### Estimated CD14+ Monocyte percentages associates with BCL-2 protein abundance

Next, we compared the effects of *BCL-2* gene expression and CD14+ Monocyte percentages with *BCL-2* protein abundance within each sample. For this purpose, we used our in-house produced proteomics data after batch correction (**Figure S6A, S6B**) for matched AML cases (n = 39; LUMC) to correlate the abundance of Apoptosis regulator *BCL-2* protein vs gene expression (**Figure S6C**) and CD14+ Monocyte percentages (**Figure S6D**). The data showed that *BCL-2* gene expression and CD14+ Monocyte percentages both correlated with *BCL-2* protein abundance to a similar extent (*R*^2^= 0.45, P < 0.001). We also overlaid plots for *BCL-2* expression and CD14+ Monocyte percentages with information on patients’ genetic abnormalities (**Figure 4A**). As expected, based on their low AUC for Venetoclax response (**Figure 3B**), two AML cases with PML-RARA had high *BCL-2* protein expression (n=2, dark green). We also confirmed the strong variability in AUC for Venetoclax response within the group of AML with mutated NPM1 (n=18, green) by showing two distinct groups at the extremes of CD14+ Monocyte percentages which correlated with *BCL-2* protein expression (**Figure 4B**). Within the group of AML patients with mutated NPM1, CD14+ Monocyte percentage associated stronger with both *BCL-2* protein abundance (R^2^= 0.56, P < 0.001) than *BCL-2* gene expression (R^2^= 0.52) (**Figure 4C, 4D**). In conclusion, the data demonstrate the relevance of our deconvolution approach on bulk RNA-Seq to separate AML, especially those with mutated NPM1, with high and low monocyte percentages to predict the patient’s response to Venetoclax.

**Figure 4.**
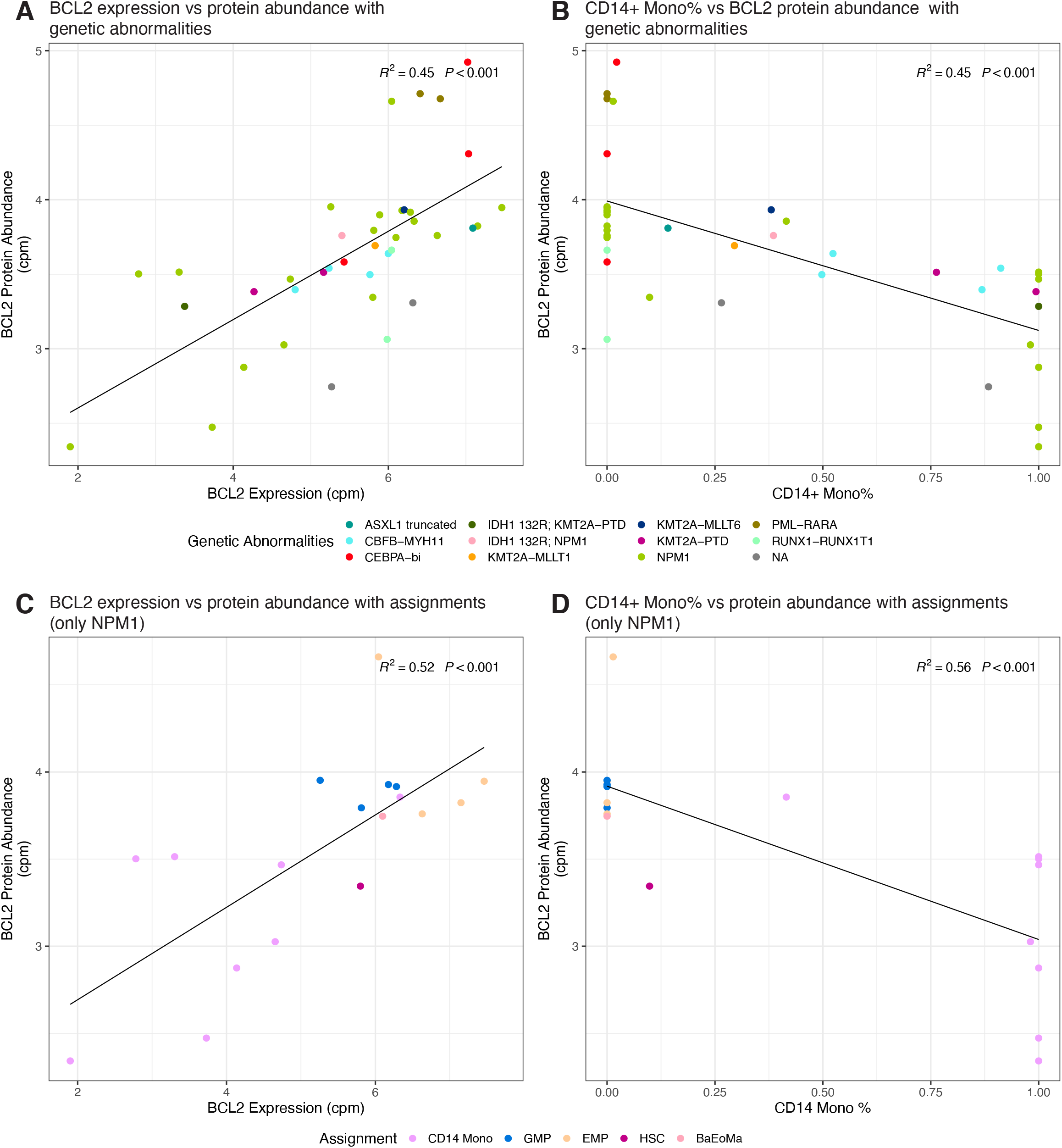
Proteomics data and ECC relations. **A** *BCL-2* expression vs protein abundance (n = 39) annotated with genetic abnormalities (R^2^= 0.45, P < 0.001). Each dot represents a sample. **B** CD14+ Monocytes percentage vs *BCL-2* protein abundance annotated with genetic abnormalities (R^2^= 0.45, P < 0.001). **C** Similar to A; but only limited to NPM1 mutated samples (n = 18) and annotated with patients’ AML phenotypes (most abundant cell type) (R^2^ = 0.52, P < 0.001) **D** Similar to B; but only limited to NPM1 mutated samples (n = 18) and annotated with patients’ AML phenotypes (most abundant cell type) (R^2^ = 0.56, P < 0.001).

## Discussion

In this work, we utilized single cell guided deconvolution to decompose bulk transcriptomics data from five independent AML cohorts and show that the obtained estimated cell compositions (ECCs) faithfully reconstitute the FAB landscape (M0-M7). Moreover, using same-sample flow cytometry, we were able to validate our deconvolution framework. Hence, following our previous work using deep transcriptomics to call various types of leukemia-defining genetic aberrations^10,26^, our current findings further underpin the power of transcriptomic-based approaches as a comprehensive and versatile platform for AML diagnosis. We next illustrated the potential use of our deconvolution framework for precision medicine applications, by correlating the estimated ECC to the results of an ex *vivo* drug resistance screening of 122 small-molecule inhibitors in the BEAT-AML study. For the *BCL-2* inhibitor Venetoclax we show that higher levels of the estimated ECC subset ‘CD14+ Monocyte’ correspond to a higher resistance, and intriguingly, that estimated CD14+ Monocyte levels is a better explanatory variable of resistance to Venetoclax than *BCL-2* expression alone. Nevertheless, using same-sample in-house proteomics data in 39 patients, we show that the estimated CD14+ Monocyte levels accurately mark BCL protein expression, and that for *NPM1*-mutated patients the presence or absence of a CD14 monocytic outgrowth corresponds with a distinct NPM1 protein abundance.

As Venetoclax was approved by the US Food and Drug Administration (FDA) in October 2020 for its use in clinics for untreated AML patients^27^, other researchers also experimented with it and also observed a similar phenomenon that we had observed. For instance, Kuusanmaki *et al.* reported that monocytic differentiation of AML reduced sensitivity to Venetoclax ex vivo^8^. Similarly, Pei *et al.* reported that the different monocytic subclones in vivo created resistance to Venetoclax treatment^9^, and also recently, White *et al.* showed that *BCL-2* inhibitor resistance could be predicted via the genes associated with monocytes. Recently, using the flow cytometry data from untreated MDS patients Ganan-Gomez et *al.* showed that after relapse and becoming secondary AML (sAML) patients, those patients with less maturated cell types (EMP) before treatment had a faster complete remission (CR) and longer relapse free status compared to more matured cell type (GMP) with Venetoclax treatment, supporting our hypothesis that ECCs could also predict Venetoclax resistance for MDS and sAML patients^28^. Collectively, these studies suggest that Venetoclax has different resistances at different maturational stages, and especially higher resistance for patients with CD14+ Monocyte dominated phenotype as we have shown in this manuscript, and we provide an open-source framework, *seAMLess,* for replicating our results or applying it to other clinically relevant datasets.

Our findings may potentially have important implications on drug use especially for genetically uncharacterized patients (AML, NOS) currently accounting for ~40% of all AML as well as other well-characterized patients such as NPM1 mutated samples. For instance, one interesting case report was from Korea ^29^, an elderly AML patient was treated with Venetoclax along with Azacitidine, and even though the patient had a favorable NPM1 status this treatment lead into tumor lysis syndrome (TLS) and patient ended up in ER 12 h into the treatment. Although the manuscript does not indicate the FAB status, AML cells expressed CD33, CD117 and myeloperoxidase, demonstrating a more stem cell than monocytic phenotype, thus one possible hypothesis could be the patient having a more sensitive cell types (e.g., EMP, GMP) to Venetoclax and thus causing immediate apoptosis of these cells and leading to TLS.

One limitation of our deconvolution strategy is that it cannot distinguish the cancerous cell types as it uses a healthy bone marrow as a reference. The reason we wanted to emphasize this is, for instance in **Figure 4D** there is a relapsed NPM1 patient with 41% of CD14+ Monocytes, as deconvolution percentage might not be exact, this patient could potentially have a healthy monocytic population but rather the reason for his relapse could be due to a different cell type, possibly 17% EMP as it is the second most abundant cell type for this patient. However, we have two rationales behind not using a single cell reference with cancerous cells. First, without mutation calling for all cells, one cannot be sure whether a cell is cancerous or not. Strategies like predicting cancer cells based on their transcriptional similarity of cells with mutation calling, proposed by Van Galen et al.^16^, adds another level of ambiguity to already not perfect deconvolution pipelines. Secondly and more importantly, heterogeneity of AML causes further sub-clustering within individual AML cases (e.g. UMAP plots of Triana *et al.,* 2021), therefore creating a not well-characterized cell type signature but rather patient specific clusters^17^. Additionally, using healthy subsets as reference allows our framework to provide more interpretable and intuitive results for clinicians and doctors, as it reports immune phenotypes and percentages on contrary to score-based prognostic values^15,24,26^. To summarize, we believe our proposed pipeline could be a blueprint for assessing new drugs’ resistances on different cell types of AML and along with our framework, they may provide better insights for clinicians and help paving the way into precision medicine in AML.

## Methodology

### Creating the healthy BM single-cell reference

We downloaded three different healthy BM datasets from two different studies, namely Van Galen et. al.^16^(full-transcriptome, n=6,915), and Triana et. al.^17^ (full-transcriptome, n = 13,165; 462 targeted mRNA, n=49,057). Then, all cell labels were uplifted up via *Seurat* package^30^ (v4.0.3) default query annotation pipeline to match with the Triana’s full-transcriptome cell labels as it had the most recent and detailed labels. Next, *Seurat’s* integration pipeline with *CCA* was run and the cells that were labeled as doublets/multiplets were removed from the down-stream analyses, and this yielded a healthy BM atlas of 69,130 cells in total, covering 439 genes. Also, we used the 40,000 cell subset of HCA provided by *SeuratObject* package (v4.0.2) for the validation analyses (**Figure 1C,1D**).

### Different schemes of creating pseudobulks

To create a cancerous-like pseudobulk profile from healthy BM reference and HCA subset, we selected a total of 1,000 cells for each profile, majority (80%) of them coming from a cell-type (over-abundant) and the rest of the cells were distributed according to inverse proportion of the numbers of cells for the remaining cell types. To achieve this, first integrated counts were exponentiated to make them non-log scale and *slice_sample* function from *dplyr* package (v1.0.7) was used with a replacement option. Then, these non-log scale cell counts were summed up to create the pseudobulk profiles. For individual-based pseudobulk, *AggregateExpression* from *Seurat* package was used.

### Flow Cytometry data

AML cases were stained with fluorescent antibodies and analyzed by flow cytometry for diagnosis, prognosis, and disease monitoring of AML in the diagnostic laboratory of the department of Hematology in the Leiden University Medical Center. The flow cytometric test has been developed and performed according to EuroFlow standard operating procedures (www.euroflow.org)^19^. EuroFlow antibody combinations have been tested against references databases of normal cells from healthy individuals and allow multidimensional identification and distinction of aberrant cells from normal cell populations. The flow cytometric test includes 8 tubes with different antibody combinations, i.e. one ALOT tube (acute leukemia orientation tube) and 7 AML tubes (AML1-7). The ALOT tube contains antibodies against CD3, CD45, MPO, CD79, CD19 and CD7. The AML tubes contain antibodies against CD16, CD13, CD34, CD117, CD11b, CD10, HLA-DR and CD45 (AML-1), CD35, CD64, CD34, CD117, CD300e/IREM2, CD14, HLA-DR and CD45 (AML-2), CD36, CD105, CD34, CD117, CD33, CD71, HLA-DR and CD45 (AML-3), NuTdT, CD56, CD34, CD117, CD7, CD19, HLA-DR and CD45 (AML-4), CD15, NG2, CD34, CD117, CD22, CD38, HLA-DR and CD45 (AML-5), CD42a and CD61, CD203c, CD34, CD117, CD123, CD4, HLA-DR and CD45 (AML-6). AML tube 7 contains antibodies against CD41, CD25, CD34, CD117, CD42b, CD9, HLA-DR and CD45, but this tube has not been used to stain AML cases analyzed in this study.

### Obtaining bulk transcriptomic data and their preprocessing

We have downloaded the non-normalized count matrices (htseq-counts) and the meta files of the four discovery cohorts (TCGA-LAML, BEAT-AML, TARGET-AML and TARGET-ALL) from https://portal.gdc.cancer.gov. For LEUCEGENE, count data was downloaded from their dedicated site (https://data.leucegene.iric.ca/)along with their provided meta data. All meta/count data were pre-processed using *R* (v4.1.0). For the meta data, genomic aberration labels were relabeled to the main AML WHO 2016 classes, non-AML samples were removed from the down-stream analyses, ELN-classes were relabeled according to ELN 2017 recommendations. For the count data, ERCC spike-ins and mitochondrial genes were removed, and the count matrix was then sorted according genes standard deviation in order to remove the duplicated genes that had less variation thus providing less information, and lastly the gene ensembl ids were converted to gene symbols. Before converting ensembl ids into gene symbols, the stemness score for each patient was calculated via count-per-million (cpm) normalized libraries, and these libraries were normalized using *cpm* function from *edgeR* package^31^ (v.3.34.1).

Our 100 AML samples (LUMC) deposited to EGA with accession number EGAS00001003096 and they are accessible upon request. QC benchmark analyses for these samples were done in our previous paper^10^. Therefore, we ran default *HT-SEQ* pipeline (v0.11.2) with paired-end option aligning fastq files to *hg38* to obtain the count matrix. All above mentioned preprocessing steps (filtering, gene name conversion) were also conducted for these samples as well before deconvolution.

### Deconvolution pipeline

To deconvolute the simulated pseudobulks and bulk RNA-seq AML patients, we used *MuSiC*^12^ package (v0.1.1) as it benchmarked highly and consistently across different cell types at various settings^11^ and had an easy-to-use open-source (GPL-3) implementation (https://github.com/xuranw/MuSiC). We used non-log scaled count values as inputs and set the normalization option to false. Patients were assigned to the groups (e.g., GMP, CD14+ Monocyte etc.) according to their most abundant deconvoluted ECCs. In the heatmap (**Figure 2A**), patients were re-ordered according to their ECCs within each assignment.

### UMAP of estimated cell compositions

First, to obtain reproducible results with umap plots, we set a seed to 2 as UMAP procedure involves random initialization. Then, we ran *umap* function with default parameters from *umap* package (v0.2.7) and used the first two reduced dimensions to create the plots. All related figures were plotted using *ggplot2* package (v3.3.5).

### Drug resistance predictions via Random Forest

BEAT-AML has drug resistance data for 122 small-molecule inhibitors, we downloaded these from their manuscript (Supplementary Table S5). Then, each drug response was min-ma× normalized, then matched with their available RNA-seq samples. Next, drug resistances were predicted with the deconvoluted ECCs. Random forest algorithm from *randomForest* package (v.4.6-14) with default parameters was used for the predictions. Each sample within each drug was predicted at leave-one-out cross-validation settings. Then, for each drug, Spearman ρ values were calculated between predicted and actual drug resistance values. To stratify the drugs according to their primary diagnosis of WHO classification, samples within each diagnosis are selected and then each drugs predicted, and normalized drug resistance were associated, then the correlation and p-values were calculated using *summary* function in base R.

### Venetoclax Association Analysis

First, each attribute was associated to standardized (min-ma× normalized) Venetoclax resistance from BEAT-AML study at univariate settings using *lm* function in R environment (v4.1.0) (Supplementary Table S11). Then, p-values and coefficients were calculated using *summary* function and then p-values were multiple hypothesis corrected using Benjamini-Hochberg procedure. For multivariate models, FAB classification, and ECC levels (except CD14+ Monocytes) were excluded as only 76 out of 460 samples of BEAT-AML had FAB classifications and as other ECCs are not independent of CD14+ Monocyte percentages (as the question is whether CD14+ Monocyte levels are independently predictive of Venetoclax resistance given *BCL-2* expression in the same model). Again, p-values and coefficients were calculated with *summary* function and plotted in a volcano plot (Figure 3F, Supplementary Table S12).

### Proteomics Sample preparation

Cell lysis, digestion and TMT labeling was performed as described in Paula and et al^32^. Cell lysis was performed using 5% SDS lysis buffer (100 mM Tris-HCl pH7.6) and 5 U benzonase nuclease (Thermo Scientific) with incubation at 95 °C for 4 minutes. Protein concentration was determined using Pierce BCA Gold protein assay (Thermo Fisher Scientific). 100 μg protein of each sample was then reduced with 5 mM TCEP. Reduced disulfide bonds were alkylated using 15 mM iodoacetamide. Excess iodoacetamide was quenched using 10 mM DTT. Protein lysates were precipitated using chloroform/methanol; resulting pellets were re-solubilized in 40 mM HEPES pH 8.4 and digested using TPCK treated trypsin (1:12.5 enzyme/protein ratio) overnight at 37□°C. Peptide concentration was then determined using Pierce BCA Gold protein assay.

The different samples, and reference samples, were arranged into five TMTpro 16plex sets. The peptides were labeled with TMTpro Label Reagents (Thermo Fisher Scientific) in a 1:4 ratio by mass (peptides/TMT reagents), total volume was 35 μL, for 1□h at RT. Excess TMT reagent was quenched with 5 μL 6% hydroxylamine for 15□min at RT. Samples corresponding to a TMT set were pooled and lyophilized.

Each TMT sample (80 ug) was fractionated by high pH reverse phase chromatrography on a Zorbax RRHD Eclipse Plus C18 2.1×150 mm 1.8-micron column, at 800 ul/min using an Agilent1200 binary HPLC system, equipped with a UV detector. The mobile phases were 10 mM Ambic pH 8.4 (A) and 10 mM Ambic/Acetonitrile 20/80 pH 8.4 (B). The gradient was from 2% to 90%B in 35 minutes. 20 fractions were collected in a circular fashion, i.e., collection per vial for 20 sec before moving to the next collection vial. After collection in the last vial collection is continued in the first vial. Fractions were subsequently freeze dried.

### Mass spectrometry

TMT-labeled peptide fractions were dissolved in water/formic acid (100/0.1 v/v) and analyzed by on-line C18 nanoHPLC MS/MS with a system consisting of an Ultimate3000nano gradient HPLC system (Thermo, Bremen, Germany), and an E×ploris480 mass spectrometer (Thermo) as in Rossi et al^33^. Fractions were injected onto a cartridge precolumn (300 μm × 5 mm, C18 PepMap, 5 um, 100 A, and eluted via a homemade analytical nano-HPLC column (50 cm × 75 μm; Reprosil-Pur C18-AQ 1.9 um, 120 A (Dr. Maisch, Ammerbuch, Germany)). Solvent A was water/formic acid 100/0.1 (v/v). The gradient was run from 2% to 40% solvent B (20/80/0.1 water/acetonitrile/formic acid (FA) v/v) in 120 min. The nano-HPLC column was drawn to a tip of ~10 μm and acted as the electrospray needle of the MS source. The mass spectrometer was operated in data-dependent MS/MS mode with a cycle time of 3 seconds, with the HCD collision energy at 36 V and recording of the MS2 spectrum in the orbitrap, with a quadrupole isolation width of 1.2 Da. In the master scan (MS1) the resolution was 120,000, the scan range 350-1200, at standard AGC target @maximum fill time of 50 ms. A lock mass correction on the background ion m/z=445.12003 was used. Precursors were dynamically excluded after n=1 with an exclusion duration of 45 s, and with a precursor range of 20 ppm. Charge states 2-5 were included. For MS2 the first mass was set to 110 Da, and the MS2 scan resolution was 45,000 at an AGC target of 200% @maximum fill time of 60 ms.

### Proteomics data processing and down-stream analysis

In a post-analysis process, raw data were first converted to peak lists using Proteome Discoverer version 2.4 (Thermo Electron), and submitted to the Uniprot database (Homo sapiens, 20596 entries), using Mascot v. 2.2.07 (www.matrixscience.com) for protein identification. Mascot searches were performed with 10 ppm and 0.02 Da deviation for precursor and fragment mass, respectively, and the enzyme trypsin was specified. Up to two missed cleavages were allowed. Methionine oxidation and acetyl on protein N-terminus were set as variable modifications. Carbamidomethyl on Cys and TMTpro on Lys and N-terminus were set as fixed modifications. Protein and peptide FDR were set to 1%. Normalization was on the total peptide amount. The 5 TMT-16plex analyses were normalized to each other by the bridge samples.

First, the abundance data is log-cpm transformed to stabilize variance among samples and then to ensure dealing with the technical batch effects, we ran *removeBatchEffect* function from *limma* package (v3.48.3) providing batch information. Then, we overlaid the transformed protein abundances onto a PCA plot to observe the clearance of batch related clustering. Then, the cell lines (positive-control) and controls (negative-control) were removed from the down-stream analyses, selecting only primary AML (n=39). Next, the transformed BCL-2 protein abundance was associated with the BCL-2 expression from LUMC cohort (**Figure 4A**)and CD14+ Monocyte percentage within each sample (**Figure 4B**). *R*^2^ and p-values were calculated using *stat_poly_eq* function from *ggpmisc* (v0.4.5) package and lines were drawn with *geom_smooth* function from ggplot2 package (v3.3.5) via method option set to linear model.

## Data analysis and reproducibility

All analyses performed in this study are provided with the following link: https://github.com/eonurk/lumc-sc-aml. We have also implemented an GPL-3 licensed R package at https://github.com/eonurk/seAMLess, which deconvolutes a given bulk RNA-seq count matrix to 22 cell types from our single-cell reference and predicts drug resistances via RF algorithm.

## Data Sharing Statement

For original data, please contact

## Acknowledgements

Authors want to thank Daniele Bizzarri for his fruitful critics on the project. This study was funded by a strategic investment of the Leiden University Medical Center, embedded within the Leiden Oncology Center, and executed within the Leiden Center for Computational Oncology. EvdA was funded by a personal grant of the Dutch Research Council (NWO; VENI: 09150161810095). The funding bodies had no role in the study design, the collection, analysis, and interpretation of data, the writing of the manuscript, and the decision to submit the manuscript for publication. Authors declare no competing interests.

## Authorship and conflict-of-interest statements

Conceptualization: MJR, EvdA, MG

Resources: EvdA

Methodology: EOK, EvdA, PvA, AMO

Investigation: EOK, JS, MG, EvdA, PvA

Visualization: EOK

Funding acquisition: EvdA

Project administration: EvdA

Supervision: MJR, MG, EvdA

Writing – original draft: EOK, MG, EvdA

Writing – review & editing: All authors

Authors declare no conflict of interest.

## Figure Captions

**Figure S1.**
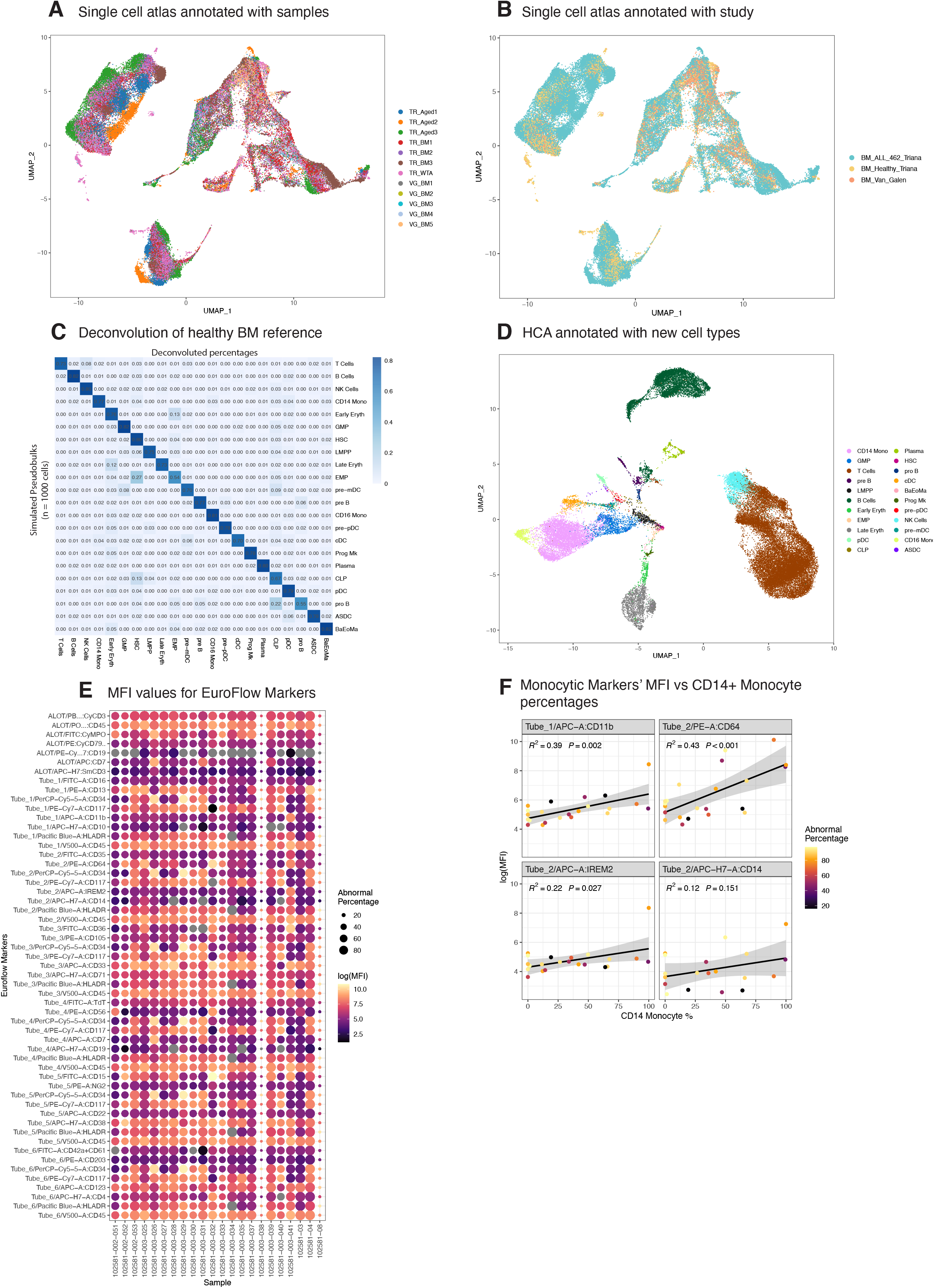
Single-cell integration evaluation & flow markers. **A** Healthy BM reference overlaid on UMAP separated by samples **B** and by different studies. **C** Similar to Figure 1D; heatmap showing the deconvolution results of simulated pseudobulks from healthy BM reference via an over-abundant cell type (80%). D UMAP for HCA subset with the cell type annotations, which were lifted via Azimuth framework. E Dot plot of MFI values of all EuroFlow markers for all samples, size of the dot represents the abnormal cell percentages per sample assigned with EuroFlow analyses. F MFI values of four EuroFlow monocytic markers (CD11b, CD64, IREM2 and CD14) and CD14+ Monocyte %. Colors indicate the abnormal percentage of cells by flow cytometry assignments.

**Figure S2.**
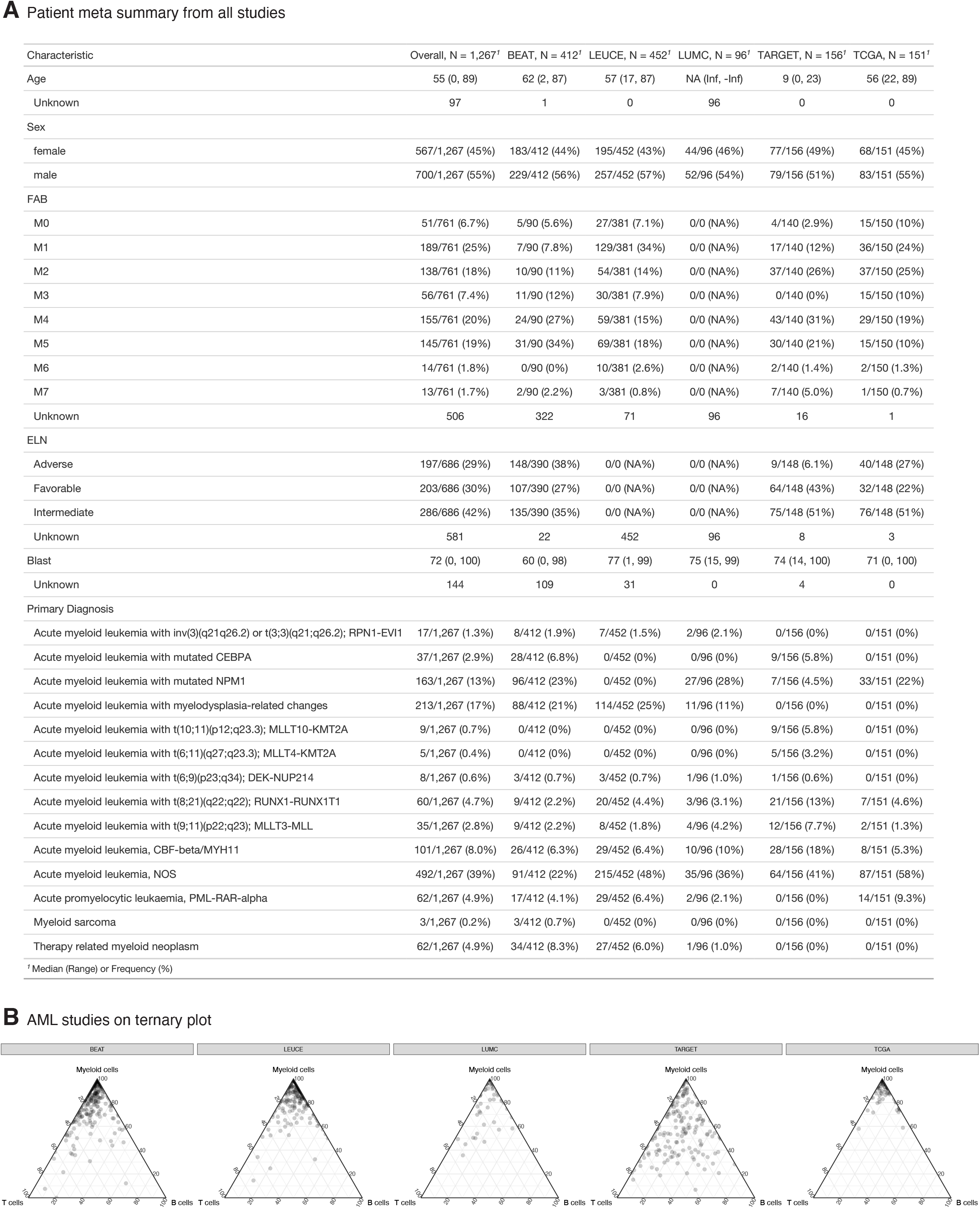
Overview of AML cohorts. **A** Detailed table for sample characteristics for age, sex, FAB, ELN, and primary diagnosis (WHO) for each cohort. Missing data is labeled as ‘Unknown’ for each attribute. *AML, NOS patients represents %39 of WHO classes. B Ternary plots like Figure 1G for all five AML cohorts.

**Figure S3.**
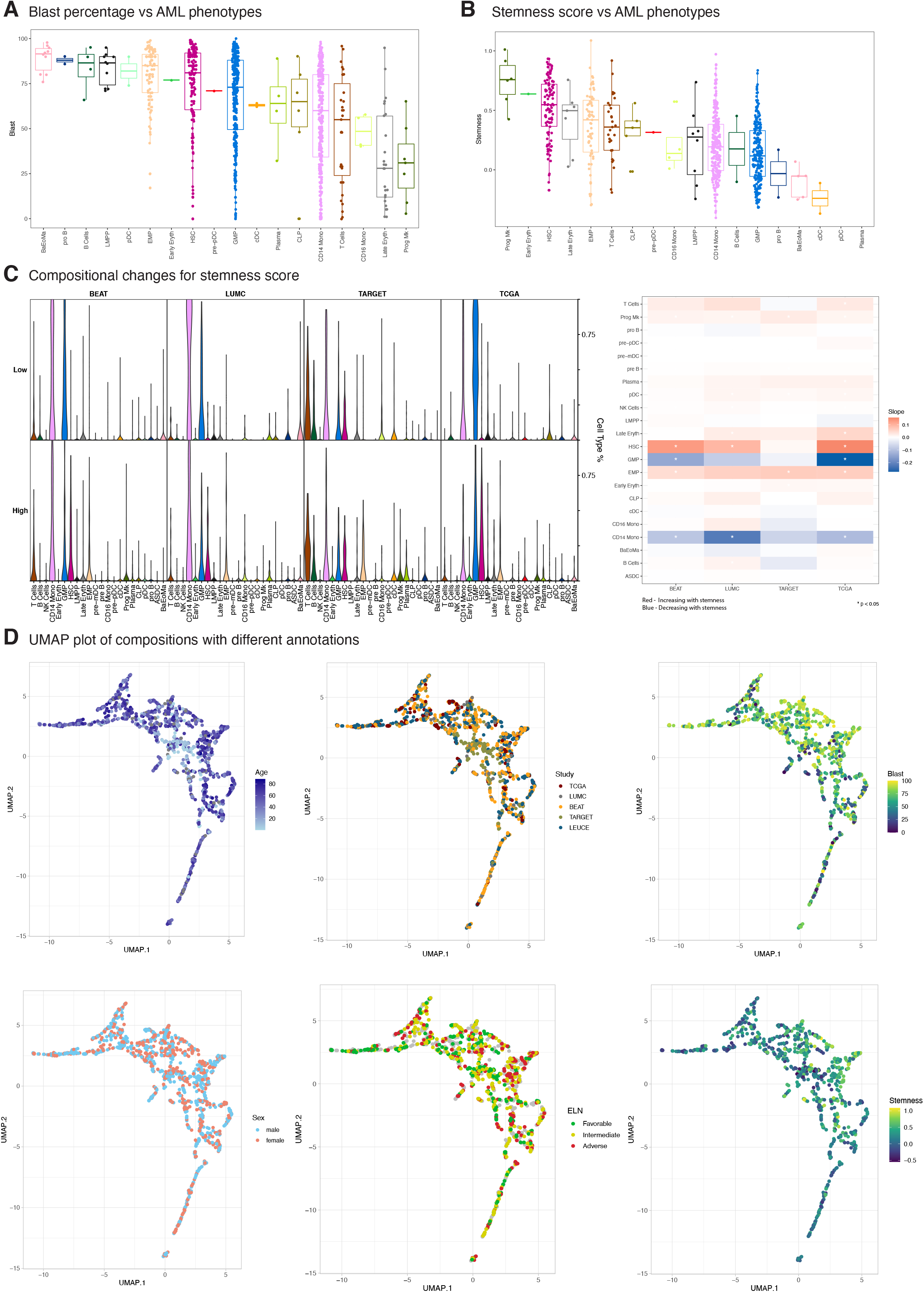
Stemness score and meta data on UMAP. **A** Box plots of blast percentage vs AML phenotype and **B** Stemness score vs AML phenotype. Boxplot were sorted according to median values. C Compositional changes of each cohort (except LEUCEGENE as it lacks entire list of genes to calculate stemness score) for low and high stemness score (split by median). Violin plots show the deconvoluted cellular percentages for 22 cell type and heatmap on the right panel summarizes the changes. A linear model is fit to calculate the slope and significance per cell type vs. stemness, and a red slope indicates more of that cell type with higher stemness score, and blue is the vice versa. Asterix indicates a statistically significant trend (P < 0.05). D UMAP plots of deconvoluted percentages (ECCs) annotated via age, study, reported blast level, sex, ELN and stemness score.

**Figure S4.**
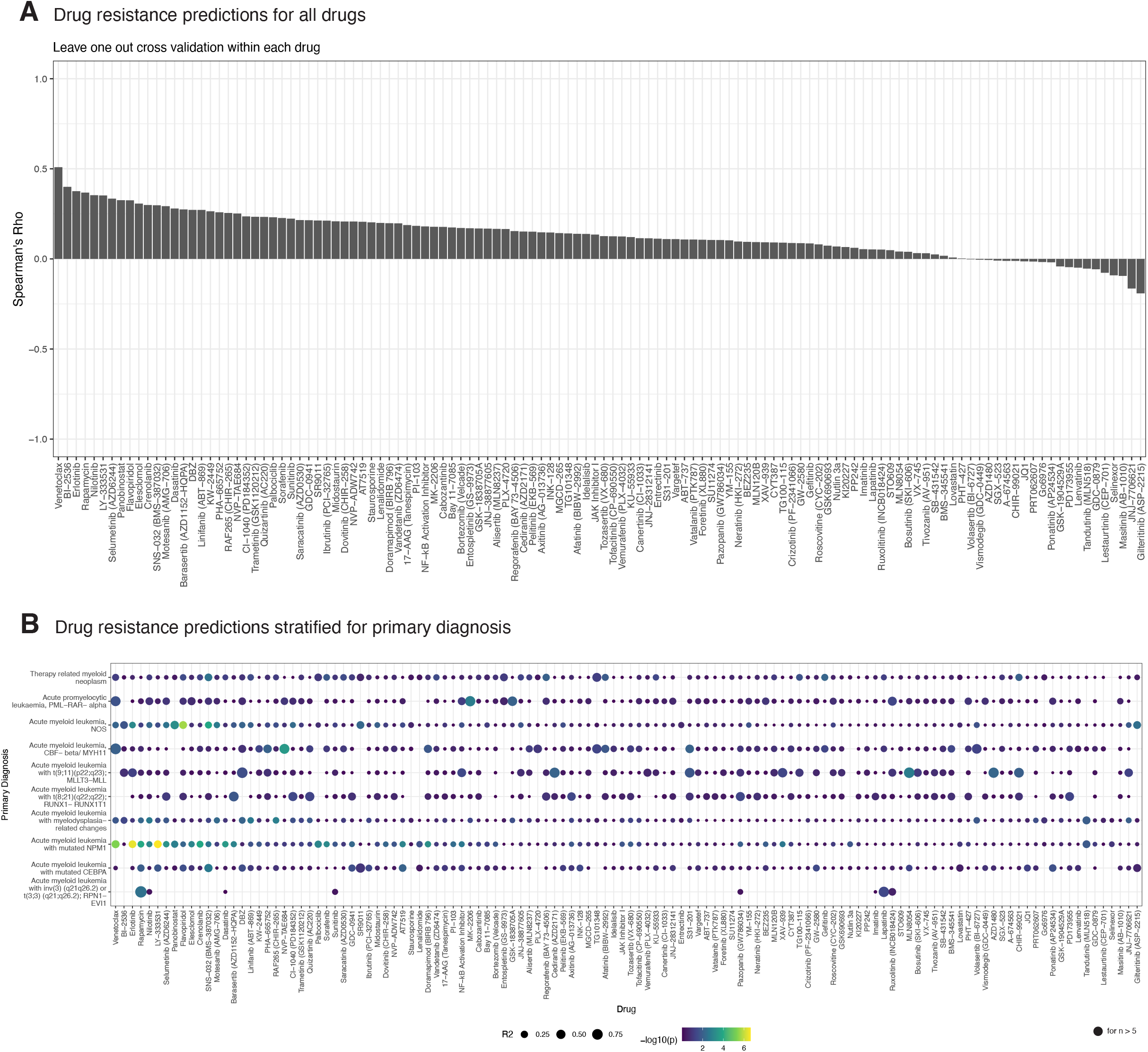
Summary plots for *ex-vivo* drug response predictions. **A** Spearman ρ values for the predictions of the drug resistance of 122 small molecule inhibitors from BEAT-AML (see Supplementary Table S7). For each drug, the resistance per sample is predicted via random forest algorithm using ECCs at leave-one-out cross validation setting. B Dot plot showing the stratified associations with predicted resistance values. Colors of the dot show the significance and sizes indicate the correlation strength. Groups with more than 5 samples were included in the plot.

**Figure S5.**
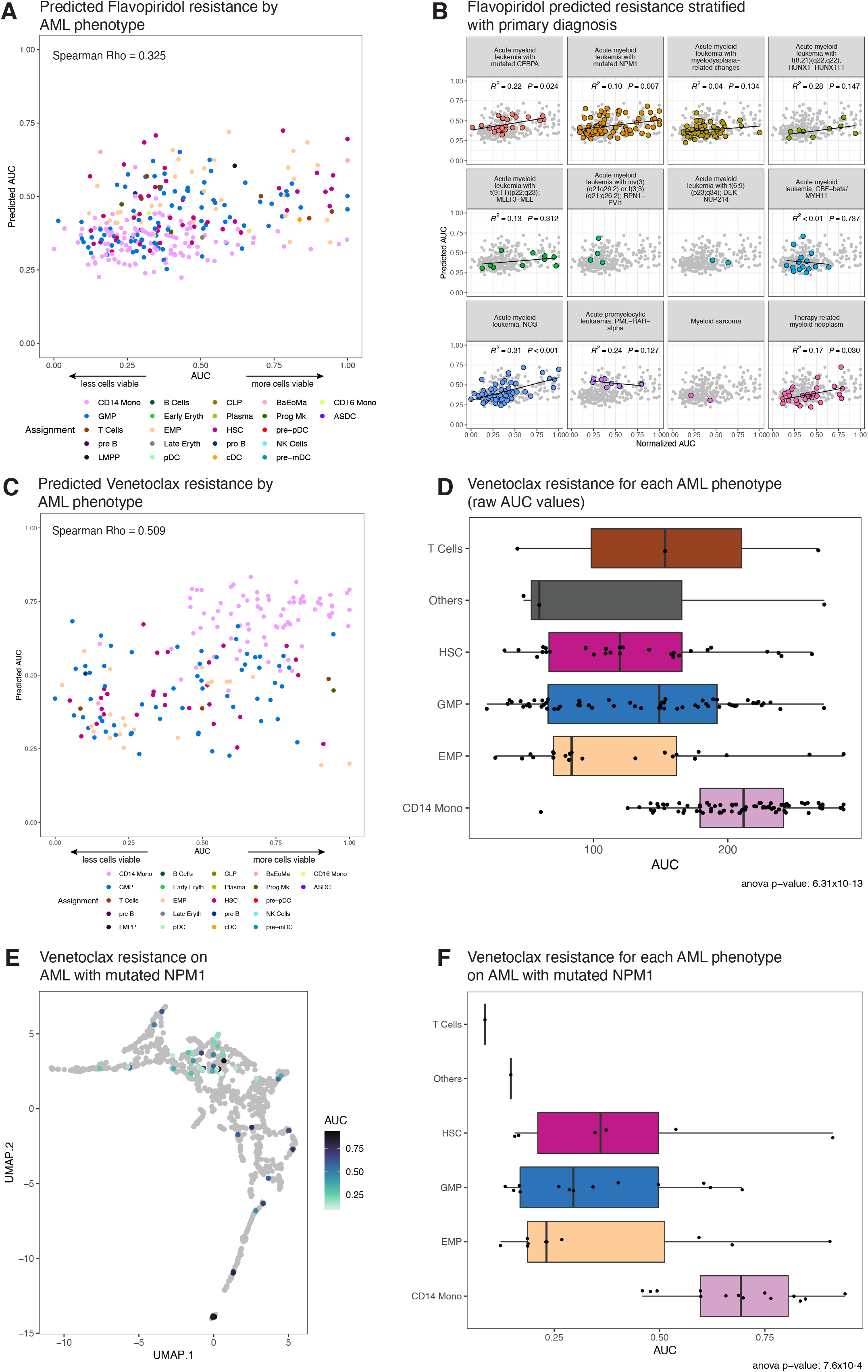
Venetoclax response on different WHO classes. **A** Flavopiridol resistance predictions with random forest annotated with patients’ AML phenotype. **B** Flavopiridol resistance predictions are faceted and highlighted by WHO classes, classes with >5 samples were fit linear models thus R^2^ and p values are shown. **C** Venetoclax resistance predictions with random forest annotated with patients’ AML phenotype. **D** Venetoclax resistance split by AML phenotypes. All drugs’ one-way anova p values are provided in Supplementary Table S9 **E** UMAP plot showing Venetoclax resistance only for NPM1 mutated samples **F** Venetoclax resistance split by AML phenotype only for NPM1 mutated samples.

**Figure S6.**
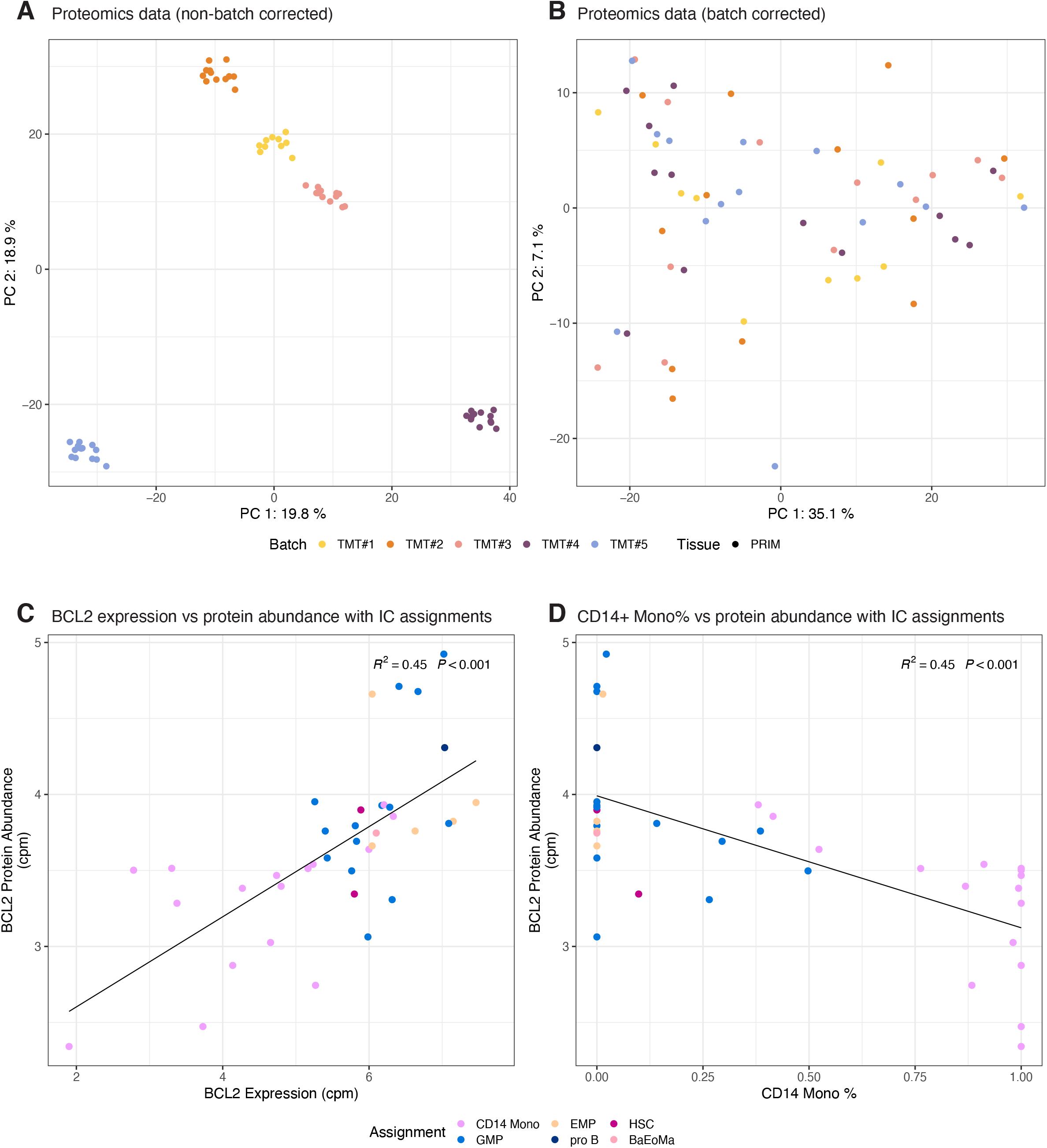
QC plots & BCL-2 expression for proteomics. **A** All-proteomics data (n = 39) without batch correction and **B** batch corrected (with *limma*) log-cpm normalized proteomics data. All abundances are log-cpm normalized. **C** *BCL-2* expression vs protein abundance annotated with patients’ AML phenotypes (R^2^= 0.45, P < 0.001). **D** CD14+ Monocytes percentage vs *BCL-2* protein abundance annotated with patients’ AML phenotypes (R^2^= 0.45, P < 0.001).

## Supplementary Tables

**Tab 1** TARGET deconvolutions

Deconvolutions of TARGET AML and ALL cohorts

**Tab 2** Deconvolution Results

Estimated cell compositions for 22 cell types along with meta data of 1,350 samples

**Tab 3** EuroFlow MFI values

EuroFlow MFI values of abnormal populations of LUMC samples

**Tab 4** EuroFlow cell type percentages

EuroFlow gating results for different cell types (abnormals are not annotated)of LUMC samples

**Tab 5** BEAT AUC

Min-max normalized AUC values for 122 small-molecule inhibitors from BEAT-AML

**Tab 6** BEAT AUC RF Predictions

Predictions of each drug resistance with random forest at LOOCV

**Tab 7** Drug Spearman Rho

Spearman rho values for drug resistance predictions of 122 small-molecule inhibitors

**Tab 8** Drug Associations stratified with WHO

Predicted drug and normalized resistance associations within each primary diagnosis

**Tab 9** Anova - Drug vs Cell Types (All)

One-way anova test results with sample phenotype assignments for drug resistances

**Tab 10** Anova - Drug vs Cell Types (NPM1)

One-way anova test results with sample phenotype assignments for drug resistances (NPM1 only)

**Tab 11** Univariate Venetoclax

The results of linear model associations of Venetoclax resistance to different attributes

**Tab 12** Multivariate Venetoclax

The results of linear model association of Venetoclax resistance with different attributes

## Notes

https://github.com/eonurk/seAMLess

### Competing Interest Statement

The authors have declared no competing interest.

### Summary of Updates

Introduction has been revised, naming has been resolved, i.e. immune composition now referred as estimated cell composition (ECC).

